# Inferring the structures of signaling motifs from paired dynamic traces of single cells

**DOI:** 10.1101/809434

**Authors:** R. A. Haggerty, Jeremy E. Purvis

**Author notes:** Corresponding Author: Jeremy Purvis, Mary Ellen Jones Building 11018C, CB#7488, 116 Manning Drive, Chapel Hill, NC 27599-7488.

## Abstract

Individual cells show variability in their signaling dynamics that often correlates with phenotypic responses, indicating that cell-to-cell variability is not merely noise but can have functional consequences. Based on this observation, we reasoned that cell-to-cell variability under the same treatment condition could be explained by a single signaling motif that deterministically maps different upstream signals into a corresponding set of downstream responses. If this assumption holds, then repeated measurements of upstream and downstream signaling dynamics in single cells could provide information about the underlying signaling motif for a given pathway, even when no prior knowledge of that motif exists. To test these two hypotheses, we developed a computer algorithm called MISC (Motif Inference from Single Cells) that infers the underlying signaling motif from paired time-series measurements from individual cells. When applied to measurements of transcription factor and reporter gene expression in the yeast stress response, MISC predicted signaling motifs that were consistent with previous mechanistic models of transcription. The ability to detect the underlying mechanism became less certain when a cell’s upstream signal was randomly paired with another cell’s downstream response, demonstrating how averaging time-series measurements across a population obscures information about the underlying signaling mechanism. In some cases, motif predictions improved as more cells were added to the analysis. These results provide evidence that mechanistic information about cellular signaling networks can be embedded within the dynamical patterns of single cells.

**Author Summary:** Cells use molecular signaling networks to translate dynamically changing stimuli into appropriate downstream responses. Specialized network structures, or motifs, allow cells to properly decode a variety of temporal input signals. In this paper, we present an algorithm that infers signaling motifs from multiple examples of an upstream signal paired with its downstream response in the same cell. We compare the predictive power of single-cell versus averaged time-series traces and the incremental benefit of adding more single-cell traces to the algorithm. We use this approach to understand how yeast respond to environmental stresses.

## Introduction

Cells interpret complex temporal patterns of molecular signals to execute appropriate downstream responses such as changes in gene expression or cell fate [1-3]. The molecular factors that participate in these signaling networks are often organized into specialized network structures, or motifs, that carry out a specific signal-processing function [4-6]. For example, a positive feedback loop can facilitate strong and irreversible responses to an upstream signal such as the commitment to cell division [7]. A negative feedback loop, such as the metabolic response to changes in blood insulin, allows cells to adapt to different levels of an upstream signal [8, 9] or to filter signaling noise [10]. More complex network motifs, such as coupled positive and negative feedback, can lead to oscillations [11, 12]. Here, we use the term “upstream signals” to refer to the inputs that initiate signaling in a particular pathway. Examples of an upstream signal include the activity or expression level of a receptor, kinase, or second messenger. These signals are decoded by specialized motifs into “downstream responses” such as changes in gene expression or epigenetic state (Figure 1A). Understanding the signaling motifs that decode upstream signals into downstream responses is a major goal of systems biology because these mechanisms define the dynamic relationships among signaling components and provide quantitative predictions about the cellular response to pharmacological intervention [13].

**Figure 1.**
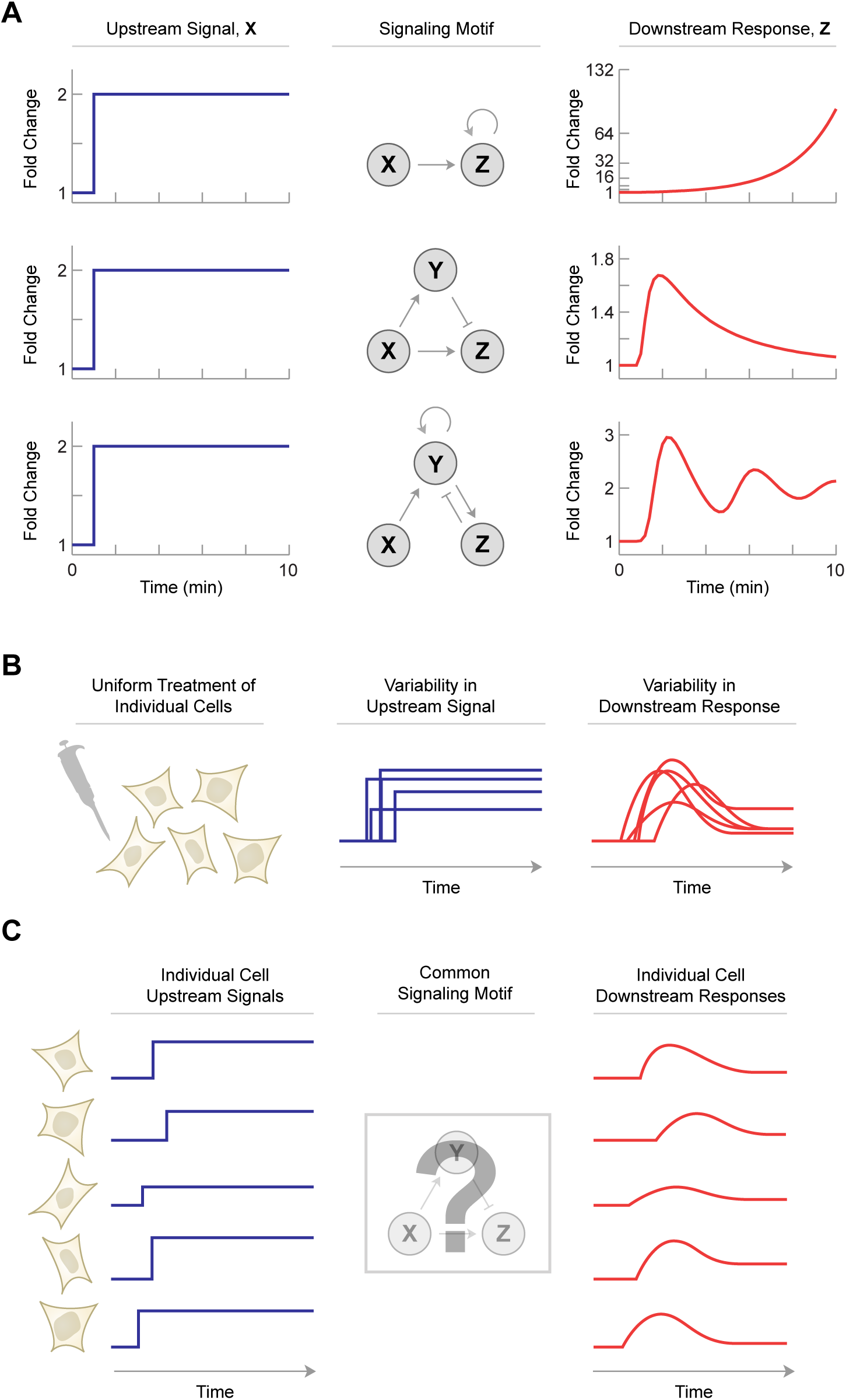
Signaling motifs determine how upstream signals are converted into downstream responses. (A) The same upstream signal, X, can produce different downstream responses, Z, depending on the signaling motif. Positive feedback leads to rapid amplification of Z following a delay in its induction. An incoherent feedforward loop (IFFL) allows Z to adapt to changes in X by first activating then dampening the downstream response. Coupled positive and negative feedback can lead to oscillations of Z. Signaling motifs often involve an intermediate signaling factor, Y, that is necessary to achieve the appropriate downstream response. Ordinary differential equations for each signaling motif are provided in the Supplemental Information. (B) In response to a given stimulus, individual cells show heterogeneous signaling patterns. For many cellular signaling pathways, the variability of the upstream signal, X, is correlated with the downstream response, Z. (C) Hypothetical model for a common signaling motif that explains the correlation between upstream signaling and downstream responses. Differences in upstream signal are mapped onto the downstream response. Under this model, it may be possible to infer the underlying structure by observing many examples of the upstream and downstream signaling patterns.

Interestingly, not all cells respond to the same upstream signal in an identical way. Previous studies have shown that individual cells show considerable heterogeneity in their dynamic responses to the same input stimulus (Figure 1B) [14, 15]. Stimulation with epidermal growth factor (EGF), for example, leads to differences in extracellular signal-related kinase (ERK2) activity [16, 17]. Similarly, uniform induction of DNA damage leads to heterogeneous patterns of p53 dynamics [18]. In many cases, these differences in signaling dynamics are correlated with distinct cell fate decisions. For example, the temporal pattern of cyclin-dependent kinase 2 (CDK2) activity after mitosis—although variable from cell to cell—predicts whether a cell will proceed directly into S-phase or spend additional time in G1 [19]. Similarly, sustained ERK2 activity following serum addition indicates that cells will enter S-phase [16]. Such correlations suggest that differences in upstream signaling patterns play an important role in driving cells toward specific downstream responses.

These collective observations suggest a testable hypothesis: if differences between individual cells have predictable effects on downstream responses, then perhaps all cells in that population share a common signaling motif that consistently interprets each cell’s unique signaling dynamics (Figure 1C). In other words, upstream signals are consistently “mapped” to downstream responses. If such a single mechanism exists in all cells, then a second hypothesis follows: by using many examples of the upstream signals and downstream responses, it may be possible to infer the underlying signaling motif without any prior knowledge of its structure. That is, single-cell signaling patterns gathered under the same experimental conditions could reveal the structure of the molecular mechanism that produced those patterns.

Here, we test these two hypotheses by analyzing paired sets of upstream and downstream signals in single cells. We focus specifically on inferring the signaling motifs that control the stress response of budding yeast [20, 21]. We introduce a computational approach called Motif Inference from Single Cells (MISC) that determines the mechanistic relationship between measurements of two fluorescent reporters in the same cell. MISC exploits the heterogeneity among identically treated cells by finding one or more signaling motifs that best explain the relationship between the paired upstream signals and downstream responses across all cells in a population. We believe this to be the first tool that infers dynamic networks from paired time-series traces from single cells. Our results provide evidence that single-cell dynamics contains information that reveals the underlying signaling mechanisms that produced them.

## Results

### Signaling motifs convert upstream signals into downstream responses

We hypothesized that cell-to-cell variation in an upstream signal can be decoded by a common signaling motif that produces an accompanying set of downstream responses. Under this model, the original source of variability in the upstream signal is not considered. Instead, we focus on the mechanistic relationship between the upstream signal and the downstream response. To explore this, we first considered how variation in an upstream signal might be propagated by a single signaling motif. We chose a simple upstream signaling pattern represented by a rapid 5-fold increase above basal levels after a 2 min delay and a return to basal levels after 6 minutes. To simulate realistic cell-to-cell variability in this signaling pattern, we allowed each cell to deviate from the average behavior in its amplitude and delay time (see Materials and Methods). This procedure allowed us to generate an arbitrarily large number of unique upstream signaling patterns that closely resembled experimental measurements from individual cells (Figure 2A).

**Figure 2.**
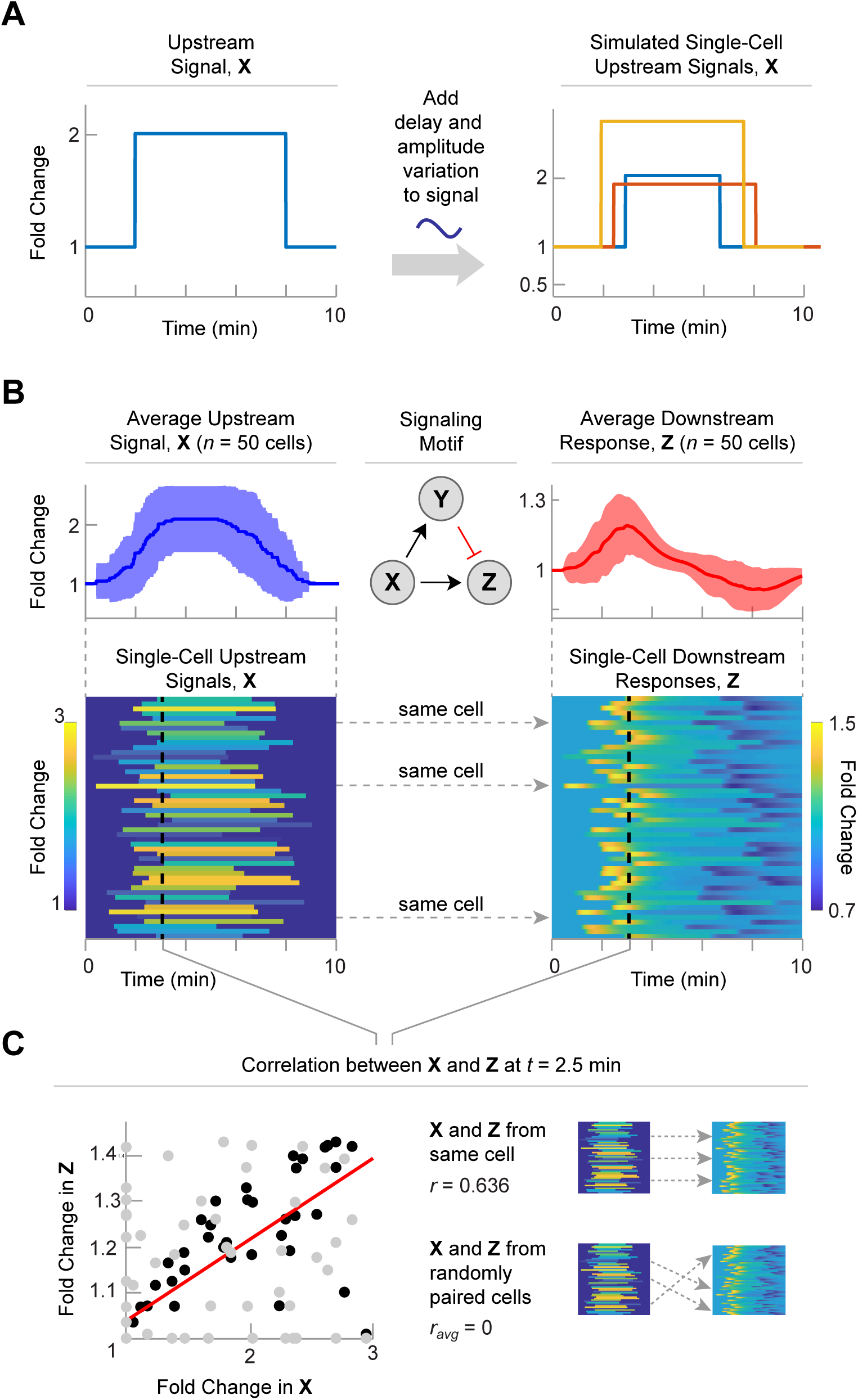
A single signaling motif can explain the correlation between upstream signals and downstream responses in single cells. (A) Experimentally measured values for cell-to-cell variation in signal height and delay were used to generate a large number of simulated signaling patterns. Each trace represents a unique upstream signal from an individual cell. (B) Average (*top*) and single-cell (*bottom*) signaling patterns for upstream signals (*left*) and downstream responses (*right*) for 50 simulated cells. While X is generated using the approach in panel A, the downstream responses are calculated using the signaling motif (middle). Here, Z if produced by the incoherent feed forward loop. A single signaling motif translates 50 upstream signals into 50 downstream responses. Each row corresponds to an individual cell. Color represents the fold change. (C) There is a correlation between the upstream and downstream signals that is not present if the cells are shuffled. Black dots are from paired cells and grey dots are from shuffled cells. The red line is a correlation line for the paired cells. The unpaired cells do not have any correlation.

We next generated 50 individual upstream signals, X, and asked how these simulated signaling patterns could be decoded by a common signaling motif (Figure 2B). We selected the incoherent feedforward loop (IFFL) as the signaling motif because of its well-characterized behavior [22, 23]. An IFFL involves three signaling factors (X, Y, and Z). X positively regulates Y and Z, and Y negatively regulates Z. The mechanism is represented as a set of ordinary differential equations (ODEs) that allow calculation of the downstream response given a particular upstream signal. We used the Hill-Langmuir equations to represent positive and negative interactions between X, Y, and Z but derived new expressions for the right hand side of the ODEs to accommodate fold-change representations of X, Y, and Z rather than absolute quantities (Supplementary Information).

As expected, the average response to the step increase in X is a transient increase in Z that returns to basal levels as the accumulation of Y dampens the increase in Z and when the signal in X disappears, the cells overcorrect and drop below basal, and then return to basal levels. At the single-cell level, each cell showed a slightly varying downstream response in accordance with the upstream signal. When the upstream signal for each cell was plotted against its downstream response at 2.5 min, these differences showed a moderately positive correlation (Figure 2C, black dots). This analysis is analogous to performing immunohistochemistry to detect levels of X and Z across a large population of cells at a fixed time point. Importantly, this correlation was destroyed when the upstream signal of a given cell was randomly paired with the downstream response of a different cell (Figure 2C, gray dots). Thus, for this analysis, the correlation between upstream signals and downstream responses among individual cells reflects a mechanistic relationship between these two signaling factors.

These results show how a single signaling motif can lead to different downstream responses by propagating variation in an upstream signal. Importantly, the signaling motif is deterministic: it does not invoke any stochastic processes to generate different downstream responses. Instead, cell-to-cell differences are each mapped to unique downstream responses.

### Inferring the signaling motif from simulated single-cell dynamics

For many signaling pathways, the signaling motif that converts upstream signals into downstream responses is unknown [16, 19, 24]. We developed a computer algorithm called Motif Inference from Single Cells, or MISC, that infers signaling motifs from measured single-cell dynamics. Before applying the algorithm to an experimental data set, we first tested MISC on the synthetic data set generated by a known signaling motif described above. Figure 3A shows the upstream signaling dynamics and downstream responses for 50 simulated cells using an IFFL as the signaling motif. To infer the signaling motif in an unbiased way, we considered all possible structures of signaling motifs that could explain the relationship between X and Z, including mechanisms that involve a potential intermediate signaling factor, Y. Although it is possible to include more than one unknown factor in our search, we limited our search to networks of 3 signaling factors as proof of principle and because the additional node, Y, can represent a subnetwork of one or more factors. We did not consider feedback to X because X is explicitly provided to the algorithm and reports on its own feedback

**Figure 3.**
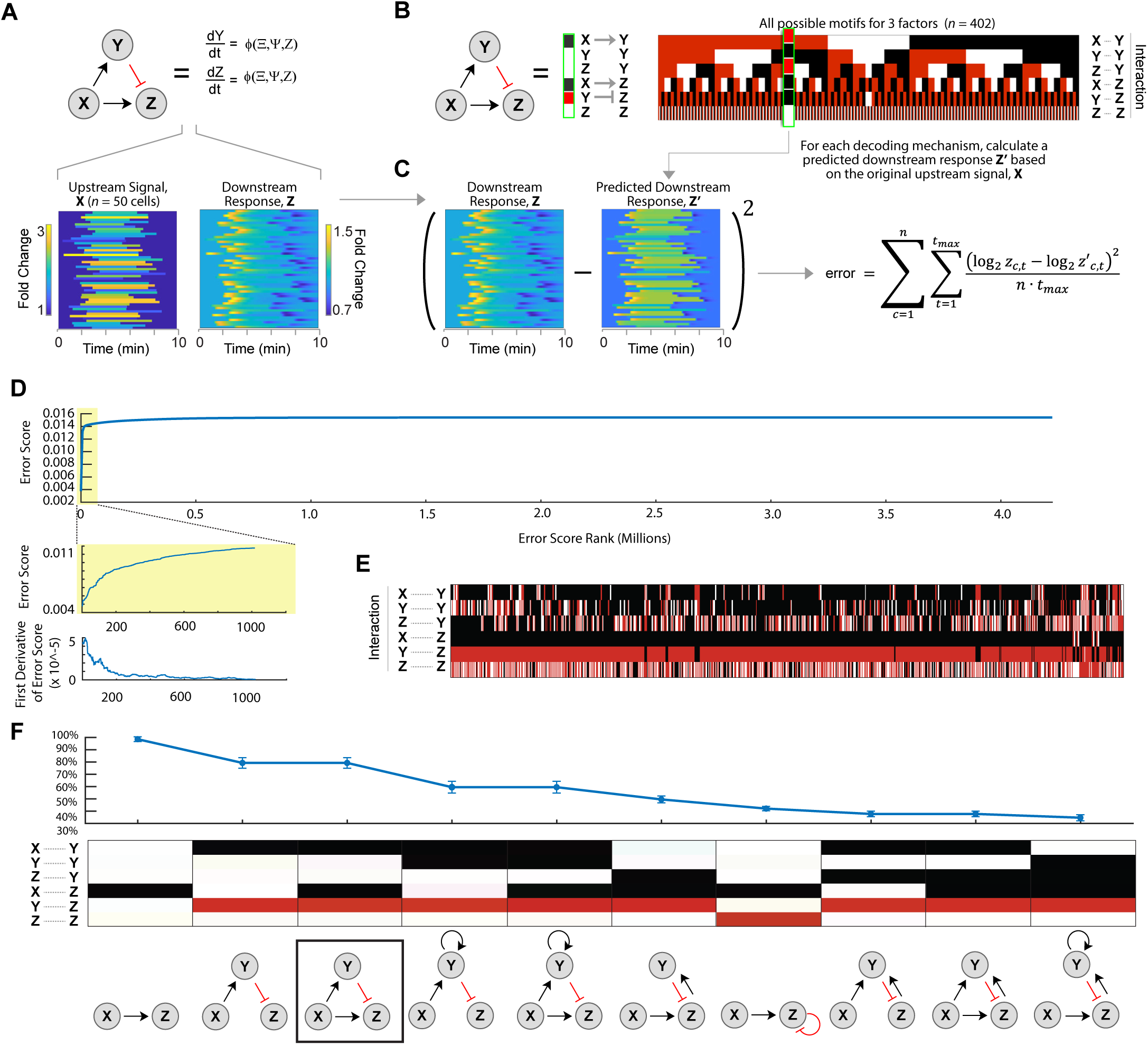
Inferring the underlying signaling motif from simulated single-cell dynamics. (A) An incoherent feedforward loop (IFFL) signaling motif was used to generate a synthetic data set for 50 single cells. Individual upstream signals X were generated as shown in Figure 2A. Each signal was then subjected to the IFFL signaling motif using ODEs to calculate a unique downstream response Z for each cell (see Supplementary Information). The resulting X and Z pairs for all cells were then analyzed to infer the underlying signaling motif. (B) Heat map representation of motif structures for all possible 3-factor signaling motifs. (*left*) A conventional representation of motif structure consisting of circles and arrows can be represented by (*right*) a single column of shaded rectangles in which each rectangle denotes a specific interaction within the motif: black, positive regulation; red, negative regulation; white, no regulation. The heat map representation for the IFFL is shown next to the conventional diagram. (Right) All 402 possible unique motif structures are shown for a 3-component network (X, Y, and Z). (C) To infer the mechanism underlying X and Z, a separate downstream response, Z’, is calculated for each possible parameterized signaling motif and compared to the original downstream response, Z. Each motif is assigned an error score that reflects the average difference in fold change between Z and Z’ per cell per unit time. (D) Distribution of error scores. The error scores are distributed such that they approach the asymptote of the worst possible score (i.e., the output of an unconnected graph). The top graphs are defined as those before the “elbow” of the graph and are determined by looking at the first derivative. (E) The top-scoring motifs. The links for the incoherent feed forward loop, as well as its simpler sub-motif structures, are overrepresented among the best-performing motifs. (F) Heat map representations of the top 10 sub-motifs (middle) and the percentage of time each motif was observed among the top-scoring results (top). Individual graphs are shown in the traditional format as well (bottom). Boxed is the IFFL, which is the top sub-motif containing all simpler sub-motifs that score better.

The complete set of signaling motifs can be succinctly represented as a heat map (Figure 3B). In this visual format, the regulatory relationships between each pair of factors is represented by a black or red box indicating positive or negative regulation, respectively. After eliminating redundant or trivial motif structures (e.g., motif containing only Y), there are 402 possible signaling motifs for a 3-factor signaling motif that allows distinct positive and negative interactions. Each mechanism was converted to set of ODEs that were used to calculate the downstream response Z for a given upstream signal X (see Supplementary Information). We fixed the sign (positive or negative) of all regulatory interactions in the mechanism and used 40,000 randomly selected, bounded, parameter sets to parameterize the ODEs (see Supplementary Information). We used a Monte Carlo approach rather than fitting parameters because we want to determine which motifs are robust to random parameterization. Next, each of the dynamic traces for this calculated response, Z’, were compared to the original downstream response to calculate an error for each signaling motif (Figure 3C). Using this approach, all signaling motifs receive an error score. Comparing these errors provides the bases for inferring the underlying signaling motif: predicted responses that best matched the original downstream response were hypothesized to be more structurally similar to the original (i.e., unknown) motif.

Figure 3D shows the errors for all randomly parameterized motifs, ranked in ascending order of error when the output of each motif was compared to the synthetic data. We found that the distribution of ranked error scores rapidly approached an asymptote of the worst possible score, which represents the output of a motif with no signaling output (i.e., a flat line at Z’ = 1). This score represents the performance of a motif with no predictive power. The vast majority of the motifs produced errors that were close to the error asymptote. To identify well-performing motifs in a principled way, we then looked for the “elbow” of the graph in which the curve began to flatten out and the errors approached the asymptote. Algorithmically, we do this by looking at the first derivative and finding the point where the slope is 1% of the maximum slope (Figure 3D, *inset*). This is the location where the slope begins to level off. After separating the motifs into well-performing and poorly-performing motifs, we look among the well-performing motifs for the most overrepresented motifs. We do this by breaking down each motif into its component motifs (e.g., the IFFL contains a 1-edge motif represented by X activating Z) and calculating the enrichment of each sub-motif among the well-performing group. Our inference approach predicted a small set of structurally similar signaling motifs including the correct motif structure. The IFFL was the top-ranking motif among those containing three interactions, present in 79% of the well-performing group. Furthermore, the top graphs also included additional motifs known to create adaptive behavior. For example, a similar structure containing positive feedback from Y to itself, was the highest ranking motif with four-interactions. Additionally, MISC predicted several motifs with negative feedback from Z to itself through Y, another motif which creates adaptive behavior in the downstream response (Figure 3F). In this manner, MISC was used to successfully infer the correct signaling motif for a synthetic data set in which the underlying motif structure was already known.

### Different yeast stress response pathways show use of the same signaling motif

We next applied MISC to determine the underlying signaling motifs from experimental data. We focused on the transcriptional response to environmental stress in budding yeast, which has been studied extensively using time-lapse fluorescence microscopy [21, 25, 26]. To predict the signaling motifs at work in the yeast stress response, we used previously published single-cell data in which two signaling activities were measured in the same cell over time [21]. Here, the upstream signal X is the transcriptional activity of the transcription factor MSN2. Upon stress, MSN2 is translocated to the nucleus where it promotes transcription of multiple target genes (Figure 4A). The downstream response Z is a fluorescent protein driven by several DNA stress response elements (STREs). Thus, X and Z represent the signaling pathway that transmits upstream stress signals to the downstream expression of target genes that allow yeast to appropriately cope with stress.

**Figure 4.**
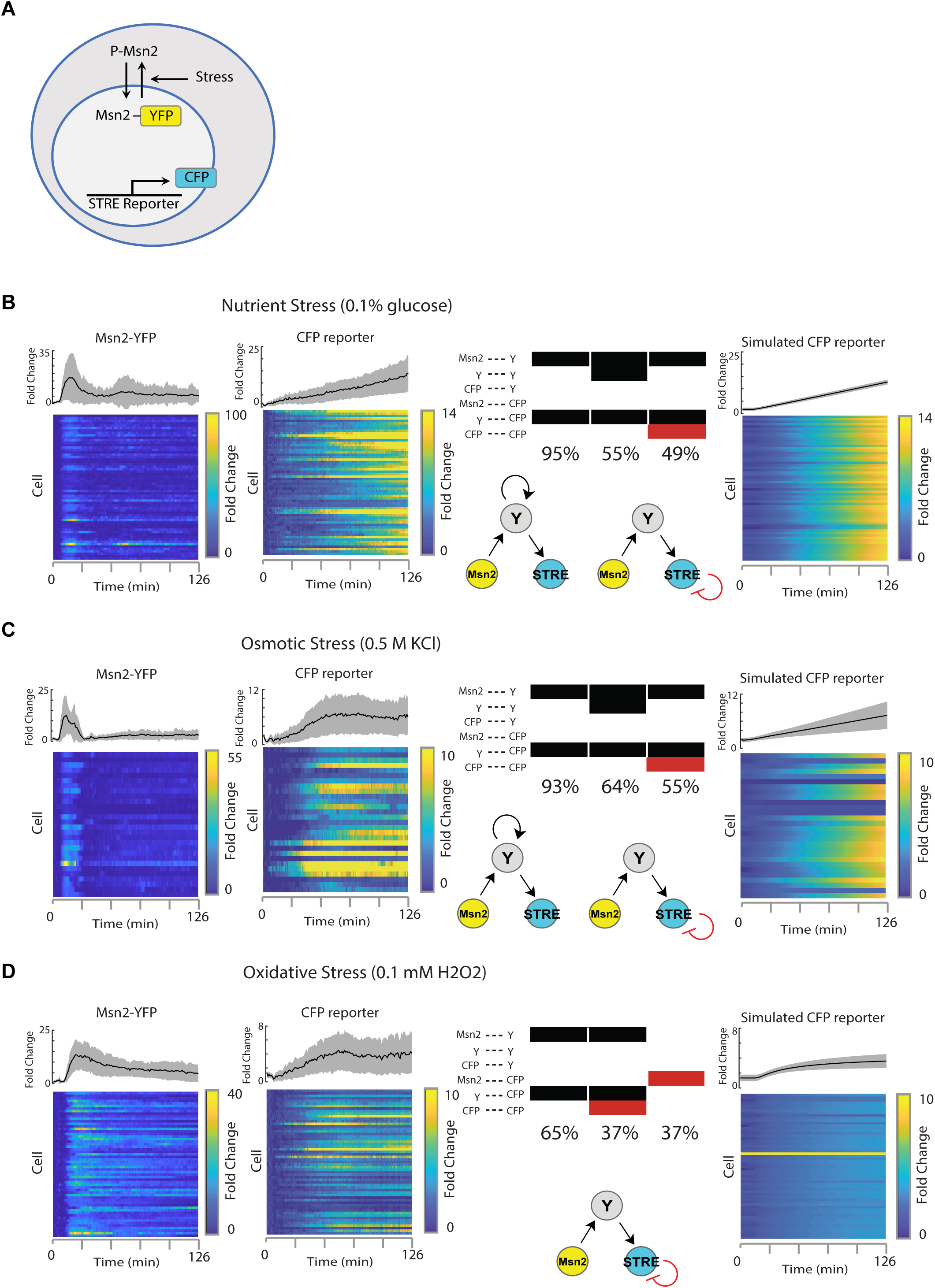
Motif inference for the yeast stress response pathway under different environmental perturbations. (A) Schematic of the system showing the fluorescent reporter system used. See ref. 21 for experimental details. Experimental data were used with author permission. (B) Oxidative Stress. 58 cells were treated with 0.1 mM H_2_O_2_. Far left, heatmap of the individual cell’s Msn2 activation, and a plot of the mean and standard deviation of the cells. Left middle, STRE response as reported by the CFP reporter. Right middle, heatmap and ball and stick representations of the significant sub-motifs. Displayed as ball and stick representations are the top sub-motifs that contain higher ranking sub-motifs. Far right, Simulated STRE response of the top preforming sub-motif. (C) Nutrient Stress. 60 cells were treated with a 0.1% glucose media. Far left, heatmap of the individual cell’s Msn2 activation, and a plot of the mean and standard deviation of the cells. Left middle, STRE response as reported by the CFP reporter. Right middle, heatmap and ball and stick representations of the significant sub-motifs. (D) Osmotic Stress. 28 cells were treated with 0.5 M KCl. Far left, heatmap of the individual cell’s Msn2 activation, and a plot of the mean and standard deviation of the cells. Left middle, STRE response as reported by the CFP reporter. Right middle, heatmap and ball and stick representations of the significant sub-motifs.

We first examined the response to nutrient stress with 0.1% sugar in 60 single cells (Figure 4B). When the activity of all cells was averaged, MSN2 showed a transient 18-fold increase followed by a return to near-basal levels. As reported previously, many cells showed a prominent pulse of MSN2 nuclear localization followed by smaller, more sporadic pulses. The averaged downstream response of STRE gene expression showed a gradual, almost linear, increase in expression. Individual cells showed considerable variability in downstream response, with some cells displaying very little changes in expression. To predict the underlying signaling motif for this stress response, we applied the MISC algorithm to the full single-cell data set. The MISC fingerprint suggested several significant motif structures that were consistent with the measured relationships between X and Z. The top ranking sub-motif was an indirect connection from X to Z mediated through Y, showing up in 95% of the top motifs. Other well performing top results included the same connection from X to Z but mediated through Y with feedback either to Y from itself (55%) or from the STRE to itself (49%). These predicted motif structures are similar to the previously published two-step model of transcription, in which the STRE promoter must first transition from an inactive to active state before productive transcription can begin [21]. Negative feedback on Z may represent first-order degradation of the transcription product, which is also present in previous models of transcription [20, 21, 26].

We next considered the yeast response to osmotic stress, using a smaller data set consisting of 28 single cells that were treated with 0.5 M KCl (Figure 4C). Under this condition, MSN2 levels showed a transient 12-fold increase in nuclear localization followed by a near-perfect adaptation to basal levels, leveling off around 3-fold. The downstream response showed a sudden increase in expression around 20 minutes after treatment that reached sustained 7-fold levels within the first 60 minutes. As with sugar stress, individual cells showed varying behavior and sporadic pulses of MSN2 activity. Applying MISC to this data set revealed the same top sub-motifs as for the glucose stress, but at slightly different percentages (93%,64% and 55%).

Finally, we considered the yeast stress response to oxidative stress using a data set of 57 individual cells (Figure 4D). Under this experimental condition, MSN2 levels showed a more prolonged response to stress, reaching 14-fold change that adapted more slowly than either salt or sugar stress. The downstream response of the STRE reporter showed a biphasic response, characterized by a gradual increase that peaked at 1 h, followed by a transient decrease and subsequent increase in expression. When these data were analyzed by MISC, the top sub-motif was the same of X to Z mediated through Y at 65%. No other sub-motifs showed up in more than 50% of the top ranking graphs, however the sub-motif with the feedback from the STRE to itself was the next highest and showed up in 37% of the graphs. Notably, the top motifs for all three stresses were similar, suggesting that even though the stresses and the responses were different, the core machinery in place to decode the stresses is the same [21].

### Single-cell dynamics contain information about the underlying motif structure that is not present in the population-averaged response

Previous studies have predicted motif structures based on a single time series [23]. It is unclear what additional value, if any, is provided by multiple single-cell measurements. To answer this question, we performed MISC on the same stress response data but permuted the data so that the upstream signal X for a given cell was randomly paired with the downstream response Z of another cell from the same experiment. Importantly, the population-averaged response remains the same under this perturbation, but the relationship between MSN2 and STRE at the level of individual cells is lost (Figure 5A).

**Figure 5.**
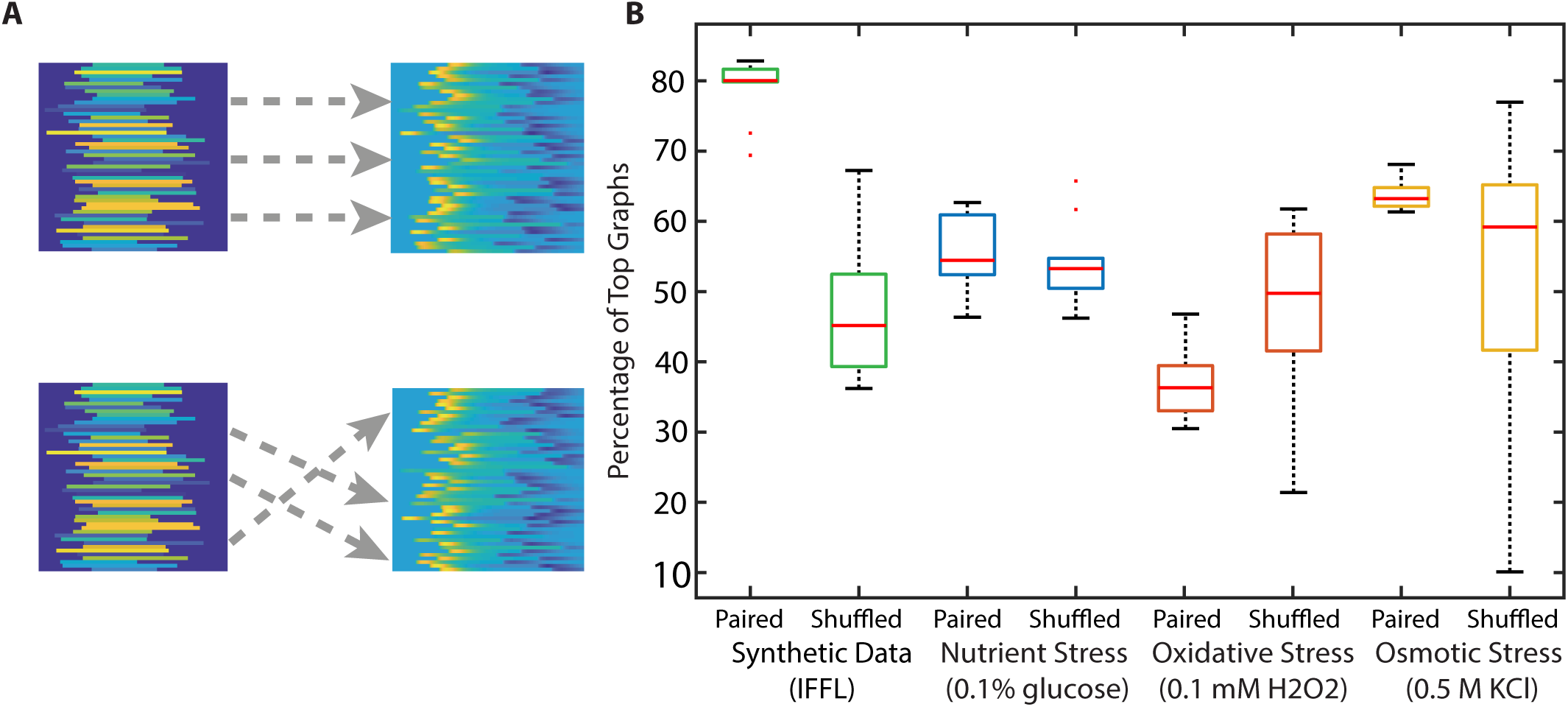
Single-cell dynamics can provide improved certainty or accuracy in motif prediction than population-averaged data. (A) In order to evaluate motif prediction based on single-cell versus population-averaged time-series traces, we performed MISC on pairs of time-series traces that were either correctly paired (*top*) or randomly shuffled (*bottom*). Note that in both cases the average time series trajectory is the same. (B) Shuffled versus paired predictions for the IFFL and yeast stresses. Ten replicates were used for each condition. Note that MISC produces variability in predictions even for paired traces since parameter values on the motifs are randomly chosen. The synthetic data, oxidative stress and osmotic stresses showed differences in distributions between shuffled and paired, while nutrient stress did not.

As proof of principal, we tested this permutation strategy on the synthetic single-cell data generated by the IFFL (Figures 2 and 3). Predictions from matched cells were consistently better at predicting the correct structure (77.5 ± 0.1%) than randomly paired cells, which predicted the correct motif in less than half of the well-performing motif structures (Figure 5B). When tested on the yeast stress response data, however, the permutation analysis showed a variety of trends. For glucose stress, random pairing of cells showed no difference in the distribution of the predictions. (KS test, p = 0.86). For both H_2_O_2_ and KCl, however, incorrect pairing of cells led to a difference in the distributions of motif predictions (KS test, p = 0.0069 and 0.0069) in which the paired cells performed better than their shuffled counterparts. For H_2_O_2_, the variance decreased when paired, and for KCl, the variance decreased and the mean increased when paired. Taken together, these results show that at least part of the predictive power of MISC comes from the observed relationships between the dynamics of X and Z in individual cells. This result suggests that information about the underlying signaling motif can be embedded in cell-to-cell variability.

We next asked whether adding additional cells to the MISC analysis could improve its predictive power. To address this question, we calculated the percentage of time the correct sub-motif appeared in the top-performing motifs as a function of the number of cells included in the analysis. As a benchmark, we subsampled the synthetic IFFL—starting with five cells—and then incrementally adding more cells until we reached the total number of cells collected under each condition. For each of these subsampled sets of cells, we ran the MISC algorithm and calculated what percentage the correct motif (as determined by using all cells) appeared among the top-scoring motifs. We performed five replicates per sample size, choosing a random subset of cells for each replicate. The IFFL showed no gain in predictive power as more cells were added to the analysis (Figure 6A). This result was expected because, in the synthetic data set, each cell’s downstream response is deterministically generated by its upstream response. The absence of intrinsic noise in the synthetic data renders each cell a perfect predictor of the underlying motif structure [29, 30].

**Figure 6.**
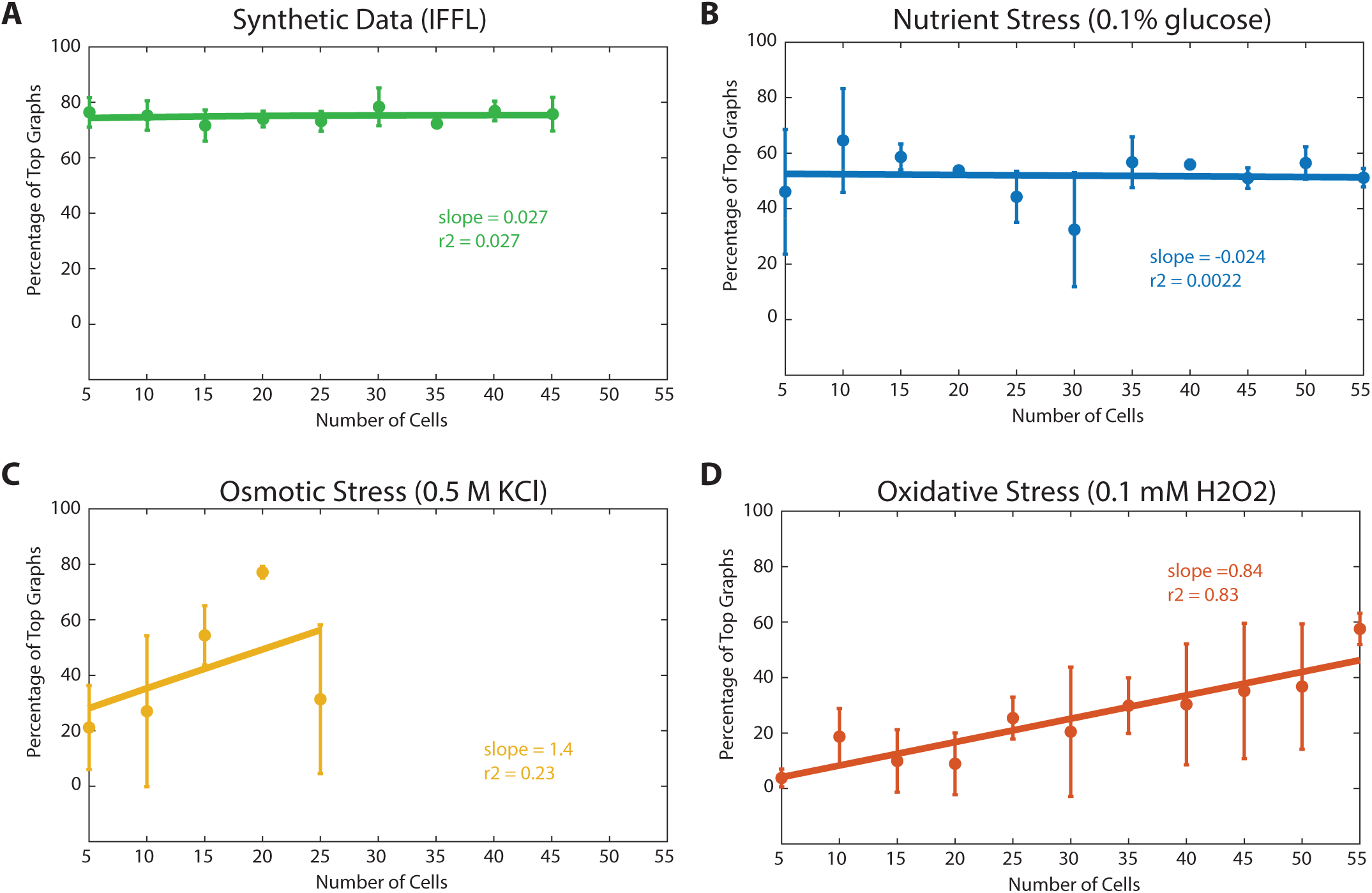
Additional single-cell measurement can provide increasingly better predictions about the underlying signaling motif. MISC was performed on an increasing number of paired single-cell measurements and the accuracy of the prediction (based on the sub-motif predicted by the full data set) was calculated. (A) Synthetic Data, (B) Nutrient Stress, (C) Osmotic Stress, (D) Oxidative Stress. The Synthetic Data and Nutrient Stress did not increase, but Osmotic and Oxidative Stresses predictions increased in percentage as the number of cells increased.

We then performed this analysis again on the experimentally-determined stress data, using the top performing sub-motif that encapsulates all higher ranking sub-motifs. Under glucose limitation, MISC also failed to show better predictive accuracy with increasing number of cells, although there was a trend toward less variance in the prediction (Figure 6B). This finding suggests that additional cells are helpful in ruling out competing models for signal transduction. Under both salt and oxidative stress, however, we observed an increase in predictive accuracy as we increased the number of single cell traces subjected to MISC (Figure 6C-D). Given the limited number of cells gathered experimentally, it is unclear at what number of cells the predictive accuracy of MISC would have leveled off. Nevertheless, these results demonstrate that each pair of time-series traces from a single cell adds incremental value to the quality of the motif prediction.

## Discussion

It is commonly stated in the field that single-cell measurements contain information that is obscured by population-averaged methods such as western blotting and quantitative PCR [31-34]. Indeed, single-cell approaches have revealed a staggering degree of heterogeneity among individual cells in terms of gene expression and protein dynamics. Here, we exploit those cell-to-cell differences by asking whether they arise from the same underlying generative process. Simulations show that we can predict a signaling motif based on paired sets of dynamic traces. We found that predictions were strengthened by considering the coupled relationships between two signals in individual cells, and that adding additional cells to the analysis can improve, in some cases, the certainty and accuracy of motif predictions. A similar strategy was used to discover signaling proteins that affect cell motility [35]. By screening a library of fluorescent fusion proteins for proteins that correlate with cell movement, this apparent “noise” in protein expression was used to identify novel regulators of motility.

Many biological network inference approaches have focused on predicting the topology of a pathway using static or population-averaged data that does not consider the single-cell dynamics of the system. Inference based on dynamical data was previously applied to global profiling data, in an effort to identify temporal transition nodes and edges [36]. MISC offers a novel inference strategy that examines a smaller scope of a biological system and aims to infer the mechanistic relationships among the measured components using single-cell dynamics. MISC allows nonlinear interactions between signaling factors. In addition, MISC allows the functional form of the governing equations to be modified to better suit various systems (Supplementary Information).

Another strength of MISC is the ability to infer components of the system that were not directly measured. Although it may be possible to measure the upstream signal and downstream response for a given signaling pathway, signaling motifs often involve additional molecular intermediates or regulatory relationships that are unmeasured or unknown. For example, it may be possible to measure the activity of a transcription factor along with target gene expression in single cells [16, 20, 37]. However, additional factors, such as cofactors or chromatin modifying complexes, may modulate the downstream response [38]. Yet, knowledge of these signaling motifs is necessary to quantify how the upstream signal is decoded to produce the observed downstream responses. We present MISC as a tool that can hypothesize unknown signaling intermediates based on the dynamical behaviors of interacting signaling factors. As with other inference methods, the motif predictions made my MISC can be experimentally tested and used to make future predictions about the cell’s behavior.

Finally, we show how increasing the number of single-cell traces improves the certainty and accuracy of motif prediction. Future studies are warranted to determine whether there is a most economical number of single-cell traces to collect under a certain experimental condition in order to identify the underlying signaling network that generated the traces. This is a practical consideration in many live-cell studies, in which accurate quantification of traces can been laborious and time-consuming, and for which there are often no best-practices for how many cells should be collected and analyzed. As such, MISC occupies a unique space in the field of biological network inference methods.

## Materials and Methods

MATLAB scripts for all simulations, calculations, and figures are provided as Supplementary Information and are available at github.com/rahaggerty/MISC.

## Acknowledgements

We thank Nan Hao and Erin O’Shea for providing raw single-cell data of the yeast stress response. We thank the laboratories of Tim Elston, Henrik Dohlman, Beverly Errede, and Galit Lahav for valuable criticism and suggested analyses. This work was supported by NIH grant DP2-HD091800 (J.E.P.) and fellowship F31-HL134336 (R.A.H.).

## Author Contributions

R.A.H. developed the MISC software, performed simulations and analysis, derived the ODE expressions for the fold-change representation of Michaelis-Menten kinetics, co-designed the figures, and co-wrote the manuscript. J.E.P. conceived of the MISC inference method, co-designed the figures, and co-wrote the manuscript.

## Supplemental Information

### Materials and Methods

#### Data Preparation

Raw data was arrays of average arbitrary fluorescent units for each cell. The first X timepoints prior to the stimulation of the cells were used as the basal levels, and the average of those timepoints was used to find the foldchange of the cells by setting those to one.

Implementation details can be found in dataprep.m

#### Parameter Bounds

a is bounded between 0 and 1 because the degradation rate must be a fraction between 0 and 1.

K, which represents the half max of the fold change is bounded between 0 and 100. This covers a broad biological range.

n is bounded between 1 and 4. The hill constant is usually between 1 and 4 for cooperative binding.

#### Coding

All code was written in MATLAB 2018a and run on UNC Research Computing’s Longleaf cluster’s general compute nodes with the following specifications:

147 compute nodes, each with 24 physical cores, 2.50 GHz Intel processors, 30M cache (Model E5-2680 v3), 256-GB RAM, and 2 ×10Gbps NIC and 30 compute nodes, each with 36 physical cores, 2.30 GHz Intel processors, 24.75M cache, 754-GB RAM, and 2 ×10Gbps NIC

### ODE Derivation

#### Hill Equations

Hill equations are widely used in biochemistry, as they provide a more accurate model of biological processes than using simple zeroth or first order kinetics. The following are the typical forms of the hill equations for activation and repression:

Activation:

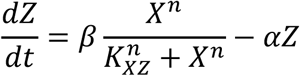

Repression:

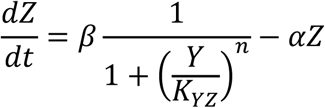

Where:

*β* = Maximal production rate of promoter

*K*_*XZ*_ = Halfway activation point of Z by X

*K*_*YZ*_ = Halfway repression point of Z by Y

*n* = Hill Coefficient, cooperativity

*α* = Degradation rate of protein

When multiple factors are affecting one node, the core of the equations are multiplied together into a compound equation to get the full effect, while the production rate and degradation rate remain the same. As an example, if Z is activated by X and repressed by Y, the total equation of the effect on Z would be:

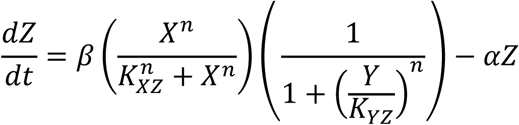

#### Autogeneration of compound equations

Given an array 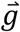 of the connections between each node to Y and Z, with one indicating activation, negative one indicating repression and zero indicating no connection, we can autogenerate the compound equations using 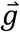 to determine which equations to include.

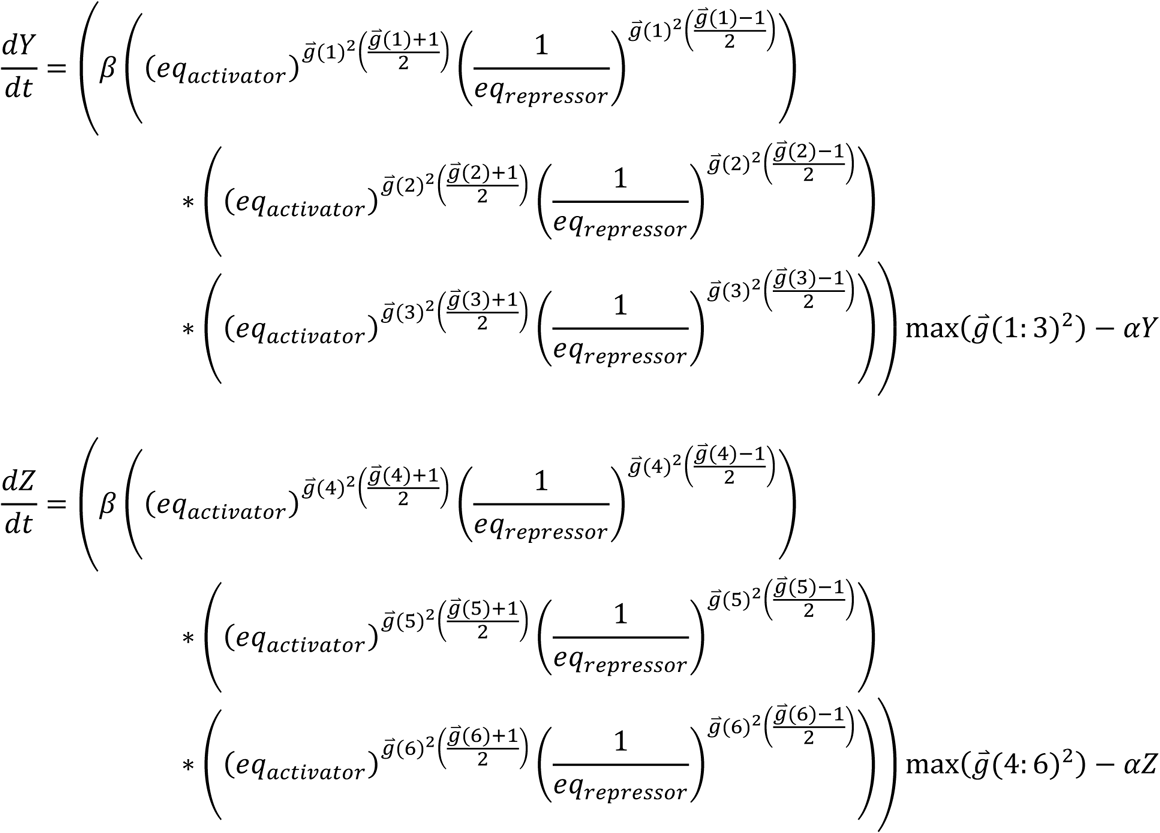

Where:

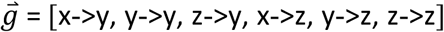

With:

0 = no connection

1 = activation

-1 = repression

And:

eq_activator_ = equation for activation of the node

eq_repressor_ = equation for the repression of the node

When the connection is 1, the activator equation is selected for that connection. If it is −1, the repressor equation is selected. If it is 0, the connection becomes one. If all connections are zero, then the selection of the maximum for the connections ensures that the total effect is zero.

#### Fold Change Equations

Hill equations are in units of concentration. However, when working with microscopy images, we are working in the realm of arbitrary fluorescent units and are unable to translate those into units of absolute concentration. We are able to get fold changes from a basal steady state. Therefore, it is necessary to get the hill equations into a form where they will work on fold change rather than absolute concentration.

At steady state, the change in Z is zero. As stated above the change in Z is as follows:

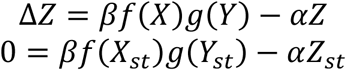

Solve for the steady state of Z:

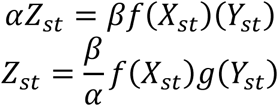

The fold change is equal to the current state of Z divided by the steady state of Z:

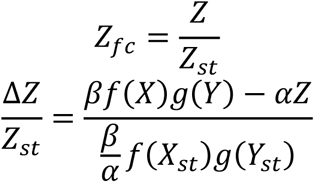

Simplify:

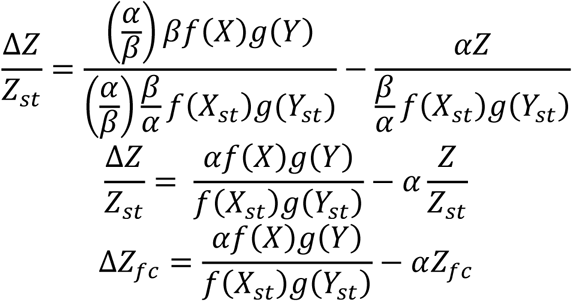

#### Full Equations

Putting together the fold change equations with the autogeneration of the compound equations, we get the following full equations:

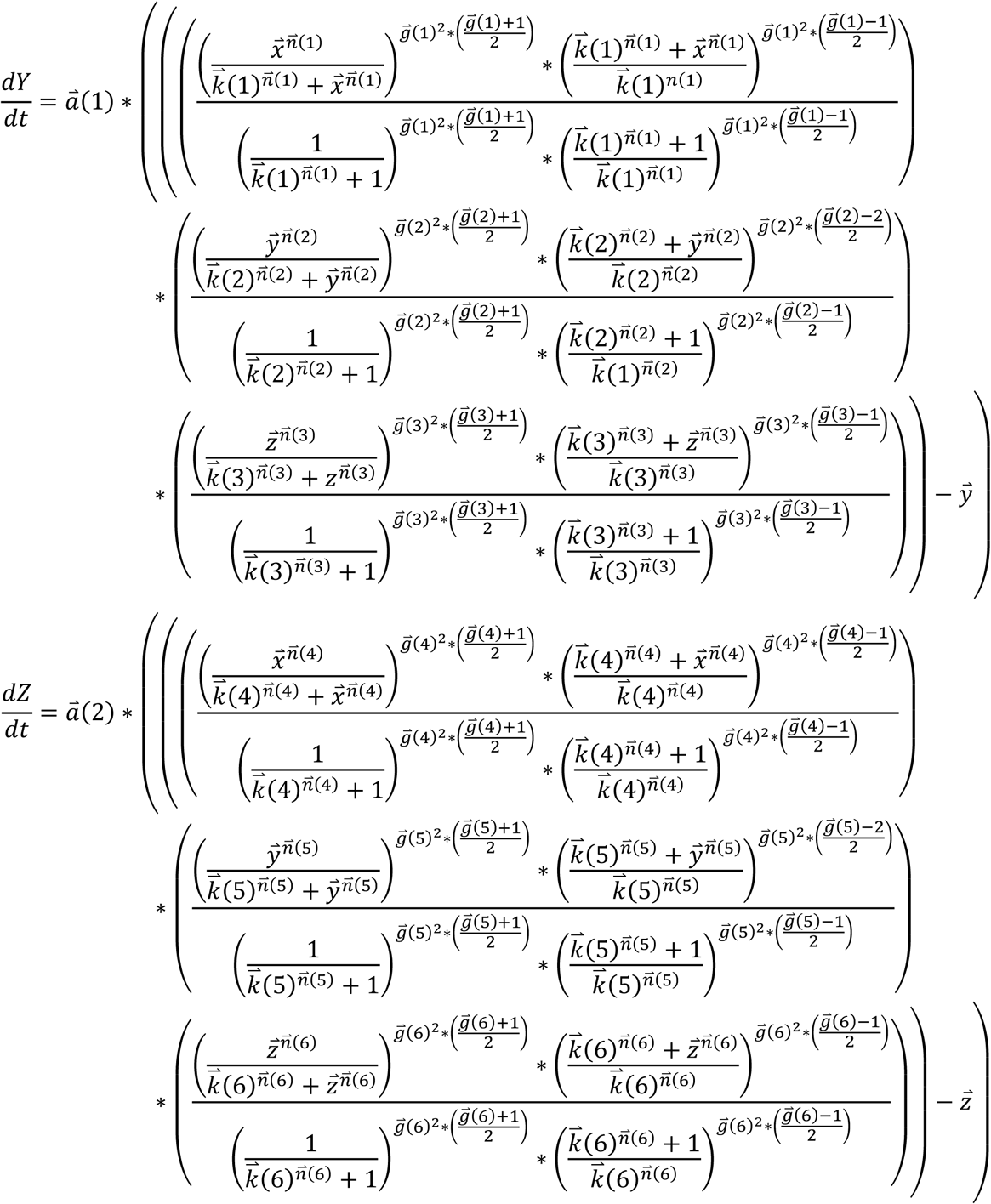

Where:

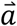 = vector with the degradation rates of y and z

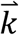 = vector of the halfway activation points

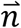 = vector of the cooperativity

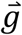 = vector of connectivity

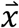 = vector of factor x

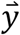 = vector of factor y

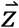 = vector of factor z

#### Code Implementation

Please see the included files dY_hill.m and dZ_hill.m to see the implementation of the full equations.

## Notes

https://github.com/rahaggerty/MISC

